# Architecture and activation of human muscle phosphorylase kinase

**DOI:** 10.1101/2023.10.22.563451

**Authors:** Xiaoke Yang, Mingqi Zhu, Xue Lu, Yuxin Wang, Junyu Xiao

## Abstract

The study of phosphorylase kinase (PhK)-regulated glycogen metabolism has contributed to the fundamental understanding of protein phosphorylation. Here we present the high-resolution cryo-electron microscopy structures of human muscle PhK. The 1.3-megadalton PhK α_4_β_4_γ_4_δ_4_ hexadecamer consists of a tetramer of tetramer, wherein four αβγδ modules are connected by the central β_4_ scaffold. The α- and β-subunits possess glucoamylase-like domains, but exhibit no detectable enzyme activities. The α-subunit serves as a bridge between the β-subunit and the γδ subcomplex, and facilitates the γ-subunit to adopt an autoinhibited state. Ca^2+^-free calmodulin (δ-subunit) binds to the γ-subunit in a compact conformation. Upon binding of Ca^2+^, a conformational change occurs, allowing for the de-inhibition of the γ-subunit through a spring-loaded mechanism. We also reveal an ADP-binding pocket in the β-subunit, which plays a role in allosterically enhancing PhK activity. These results provide unprecedented molecular insights of this important kinase complex.

## Introduction

Reversible protein phosphorylation, mediated by the interplay between >500 kinases and >100 phosphatases in human, is of central importance in physiology and disease ^1–3^. This fundamental regulatory mechanism is first discovered by Fischer and Krebs in their studies of the glycogen metabolism ^4,5^. Glycogen is the storage form of glucose in animals, and glycogen phosphorylase (GP) catalyzes the rate-limiting step in glycogenolysis to break down glycogen into glucose. Phosphorylase kinase (PhK) phosphorylates GP and transforms it from the inactive *b* form to the active *a* form to initiate this important biochemical process. The discovery of this phosphorylation-mediated regulation mechanism and the identification of PhK were major milestones in biochemistry and cell biology, as they not only provided crucial insights into the energy production process in human, but also revolutionized our understanding of cellular signaling events.

PhK is one of the largest and most complex protein kinases. It contains four subunits: α, β, γ, and δ, which assemble into a 1.3-megadalton α_4_β_4_γ_4_δ_4_ hexadecamer with a butterfly-like appearance ^6,7^. The α- and β-subunits are >1,000 residues large proteins, and together account for >80% of the molecular weight of the PhK complex ^8^. However, limited information is available regarding their 3D structures ^9,10^. The δ-subunit was identified as calmodulin ^11,12^. Unlike the interactions between calmodulin and many other calmodulin-binding proteins however, calmodulin binds tightly regardless of the presence of Ca^2+^ and serves as an integral component of the PhK holoenzyme. The γ-subunit bears the catalytic activity, and consists of an N-terminal kinase domain (KD) and a C-terminal regulatory domain (CRD). The crystal structure of the rabbit muscle PhK γ-subunit (rPhKγ) KD reveals a canonical protein kinase fold that resembles the cAMP-dependent protein kinase (PKA) ^13,14^. A significant difference is that in rPhKγ, a highly conserved Glu182 is in the place of phosphorylated Thr197 in the activation segment of PKA. This Glu182 forms an ion pair with Arg148, which comes before the catalytic Asp149. As a result, the rPhKγ-KD has a constitutively active conformation without needing to be activated by phosphorylation. The CRD contains calmodulin-binding motifs ^15^. The truncated γ-subunit without CRD is constitutively active ^16–18^, suggesting that CRD inhibits KD’s activity. Ca^2+^ activates PhK activity, presumably by binding to calmodulin and causing conformational changes; however, the molecular mechanism remains insufficiently understood.

Here we investigate the molecular architecture and activation mechanism of PhK. We determined a high-resolution cryo-electron microscopy (cryo-EM) structure of human muscle PhK in the inactive state, which elucidates the architecture and subunit organization of the PhK hexadecamer. The C-terminal region of the γ-subunit CRD docks onto the α-subunit and inhibits KD using a pseudo-substrate mechanism. Ca^2+^-free calmodulin is attached to the γ-subunit in a compact conformation, and interacts with both the N-terminal region of CRD and KD. We further studied the cryo-EM structure of PhK in the presence of Ca^2+^, and propose a spring-loaded mechanism of how Ca^2+^-induced conformational change of calmodulin de-inhibits the kinase activity. We also reveal an ADP-binding pocket in the β-subunit, and discuss the role of ADP in allosterically enhancing PhK activity.

## Results and Discussion

### Overall structure of the PhK holoenzyme

The four subunits of human muscle PhK were co-expressed in HEK293F cells, and the resulting enzyme complex was isolated with a high degree of purification (Fig. 1a, b). This recombinant PhK exhibited a robust Ca^2+^-dependent kinase activity towards GP (Fig. 1c), which is similar to that of PhK purified from rabbit skeletal muscle ^19,20^. The structure of the human muscle PhK in its inactive state, prepared in the presence of the chelating agent ethylenediaminetetraacetic acid (EDTA), was determined using the single particle cryo-EM method, with an overall resolution of 2.9 Å (Extended Data Fig. 1, Table 1). The density of one half of the complex is superior to that of the other half. Local refinements were further performed for the αβγδ and γδ subcomplexes that display good densities, resulting in 2.8 Å and 3.1 Å reconstructions, respectively (Extended Data Fig. 1a). Together, these results enable us to clearly visualize all four subunits of PhK (Extended Data Fig. 1i–l) and generate a model for the entire hexadecamer (Fig. 1d). Consistent with previous observations ^6,7^, the PhK complex exhibits a butterfly-like structure, with a central body consisting of the β_4_ homotetramer and four wings comprised of the αγδ heterotrimers. The α-subunit interacts with both the β- and γ- subunits to anchor the αγδ subcomplexes onto the main β_4_ body (Fig. 1e). The δ-subunit or calmodulin is located on the periphery of the PhK holoenzyme and interacts exclusively with the γ-subunit. We also analyzed the cryo-EM structure of active PhK in the presence of Ca^2+^ and determined the structure at a resolution of 2.9 Å (Extended Data Fig. 2, Table 1).

**Fig. 1.**
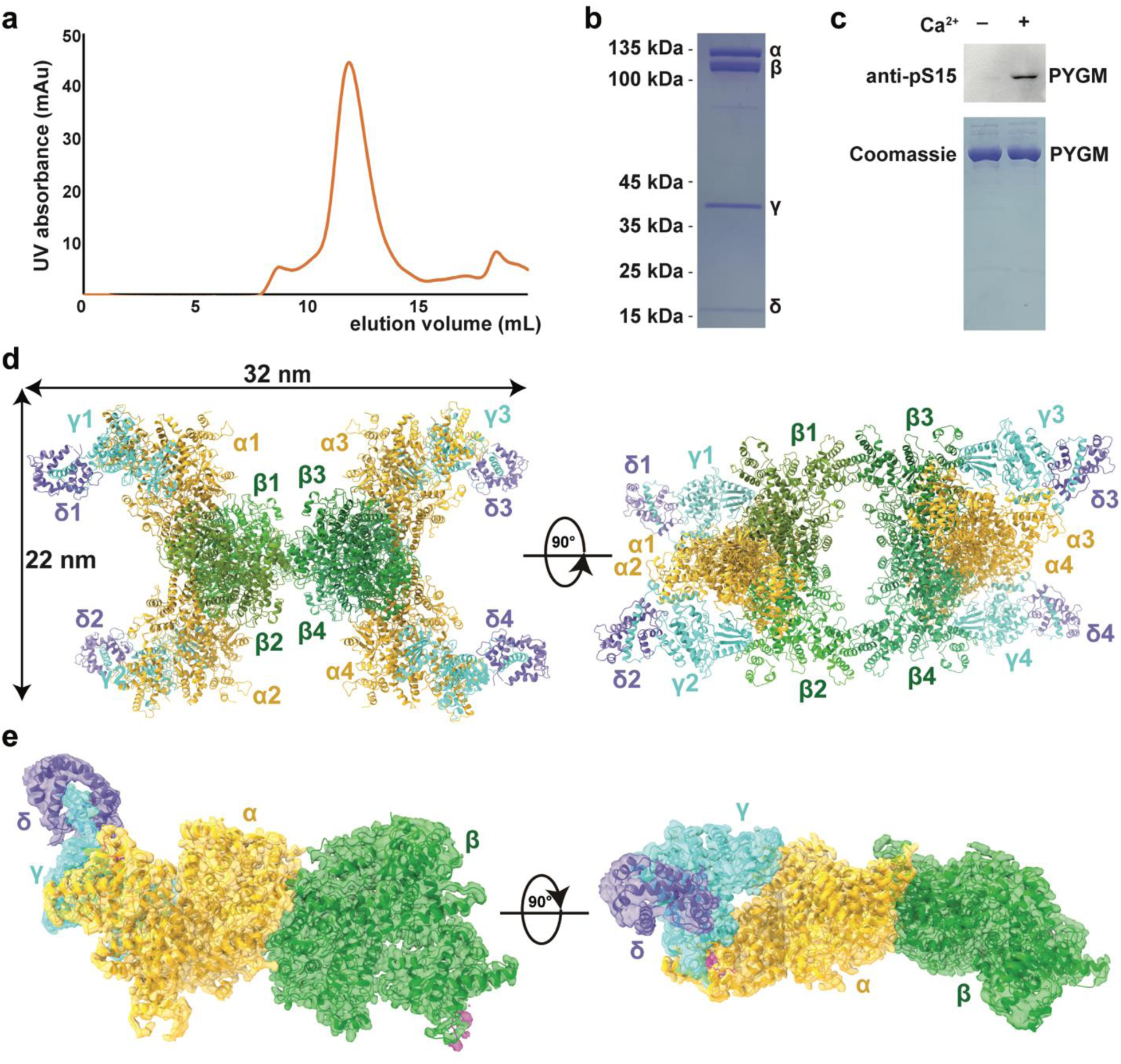
Overall structure of the PhK holoenzyme. **a.** Size-exclusion chromatography of the PhK holoenzyme. **b.** SDS–PAGE analysis of the PhK holoenzyme. **c.** PhK displays Ca^2+^-dependent kinase activity. **d.** Model of the PhK hexadecamer shown in two orientations. The β-subunits are shown in different shades of green; whereas the α-, γ-, and δ-subunits are shown in gold, cyan, and dark blue, respectively. **e.** Local cryo-EM density map of the αβγδ heterotetramer shown in two orientations, with the structural model enclosed.

**Table 1.** Cryo-EM data collection, refinement and validation statistics.

|  | Inactive_tetramer | Inactive_monomer | Inactive_GC | Active_tetramer | Active_AG |
| --- | --- | --- | --- | --- | --- |
| **Data collection and processing** |  |  |  |  |  |
| Magnification |  | 81,000 |  | 81,000 |  |
| Voltage (kV) |  | 300 |  | 300 |  |
| Electron exposure (e-/Å<sup>2</sup>) |  | 60 |  | 60 |  |
| Defocus range (µm) |  | -1.1 to -1.5 |  | -1.1 to -1.5 |  |
| Pixel size (Å) |  | 1.07 |  | 1.07 |  |
| Symmetry imposed |  | C1 |  | C1 |  |
| Initial particle images (no.) |  | 3,769,745 |  | 2,434,165 |  |
| Final particle images (no.) |  | 632,973 |  | 432,047 |  |
| Map resolution (Å) |  |  |  |  |  |
| FSC threshold: 0.143 | 2.92 | 2.72 | 3.12 | 2.94 | 2.91 |
| **Refinement** |  |  |  |  |  |
| Initial model used (PDB code) | 1CDL | 1CDL | 1CDL | N/A | N/A |
| Model resolution (Å) |  |  |  |  |  |
| FSC threshold: 0.143 | 2.92 | 2.72 | 3.12 | 2.94 | 2.91 |
| Map sharpening *B* factor (Å<sup>2</sup>) | 71.3 | 70.7 | 86.2 | 83.9 | 76.6 |
| **Model composition** |  |  |  |  |  |
| Non-hydrogen atoms | 81,204 | 20,301 | 5,480 | 65,888 | 8,091 |
| Protein residues | 10,108 | 2,527 | 679 | 8,204 | 1,018 |
| Ligands | 8 | 2 | 1 | 12 | 1 |
| ***B* factors (Å<sup>2</sup>)** |  |  |  |  |  |
| Protein | 123.99 | 56.67 | 83.39 | 96.93 | 53.55 |
| Ligand | 158.78 | 93.00 | 80.22 | 101.91 | 87.86 |
| **R.m.s. deviations** |  |  |  |  |  |
| Bond lengths (Å) | 0.006 | 0.007 | 0.006 | 0.006 | 0.005 |
| Bond angles (°) | 1.360 | 1.183 | 1.518 | 1.110 | 1.107 |
| **Validation** |  |  |  |  |  |
| MolProbity score | 2.03 | 1.90 | 1.54 | 1.95 | 1.93 |
| Clashscore | 8.78 | 8.01 | 7.87 | 7.40 | 7.52 |
| Poor rotamers (%) | 3.35 | 2.78 | 0.83 | 2.68 | 2.26 |
| **Ramachandran plot** |  |  |  |  |  |
| Favored (%) | 97.08 | 97.37 | 97.46 | 96.62 | 96.29 |
| Allowed (%) | 2.85 | 2.59 | 2.54 | 3.38 | 3.71 |
| Disallowed (%) | 0.07 | 0.04 | 0.00 | 0.00 | 0.00 |

### Structures of the α- and β-subunits

The structures of the α- and β-subunits are homologous to each other, as suggested by sequence analyses ^21^. Each can be further divided into five domains (D1–D5, Fig. 2a). The D1 domains display a glucoamylase-like fold ^22^. For example, the D1 domain of the α-subunit (D1_α_) can be superimposed to a glucoamylase-family protein with a root-mean-square deviation (rmsd) of 2.9 Å over 330 residues (Fig. 2b). Furthermore, D1_α_ appears to possess an intact active site, including the two Glu (Glu185, Glu371) that participate in catalysis. In contrast, D1_β_ lacks some of the catalytic residues ^23^. Nonetheless, both the αγδ subcomplex (AGC, Extended Data Fig. 3a) and the PhK holoenzyme do not exhibit detectable glucoamylase activity in vitro when tested against maltose, isomaltose, or glycogen (Fig. 2c). To the best of our knowledge, this is the first direct experiment conducted to investigate the glucoamylase activity of PhK. The physiological significance of these glucoamylase-like domains still requires further investigation, especially regarding D1_α_, as it appears to have a functional active site. On the other hand, it is likely that these domains are responsible for binding to acarbose, a glucoamylase inhibitor and an anti-diabetic drug that has been demonstrated to bind to PhK and enhance its kinase activity ^24^.

**Fig. 2.**
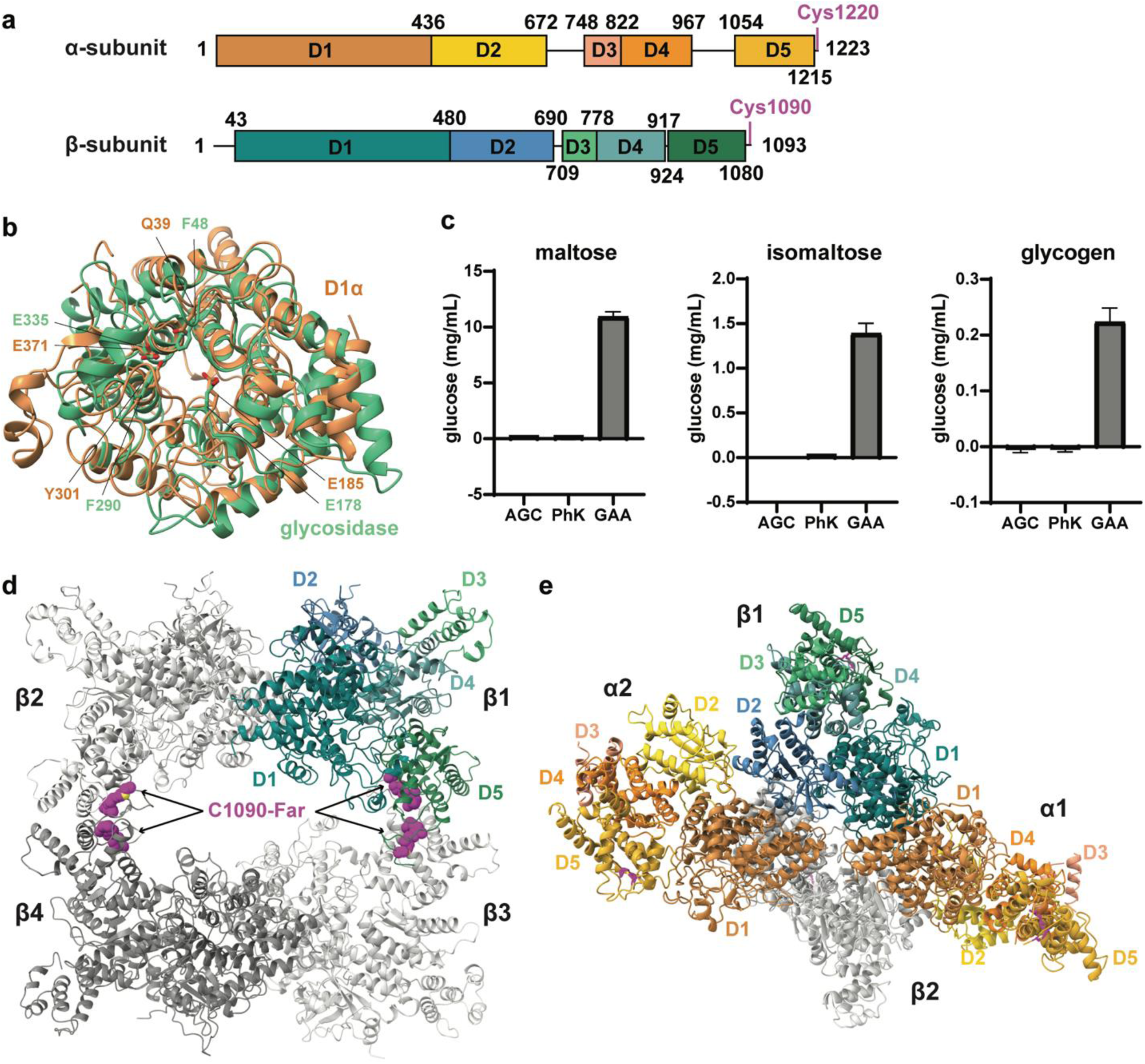
Structures of the α- and β-subunits. **a.** Schematic representations of the α- and β-subunits. **b.** Structural overlay of the D1 domain of the α-subunit and a glycosidase (PDB ID: 5Z3F), shown in yellow and green, respectively. **c.** Neither the AGC subcomplex nor the PhK holoenzyme has detectable glucoamylase activity against maltose, isomaltose, or glycogen. **d.** The β_4_ tetramer core structure. The five domains of β1 are shown in blue and green, whereas the other three β-subunits are shown in grey. The farnesyl groups are highlighted in magenta. **e.** Structure of the α2β2 heterotetramer. The five domains of the two α-subunits are shown in yellow and orange.

The D2 domains bear resemblances to a part of a glucanotransferase (Extended Data Fig. 4a), and therefore may originate from a degenerate glycoside hydrolase. The D3 domains exhibit a helical bundle structure and are the most flexible regions in both the α- and β-subunits. The D4 and D5 domains are homologous to each other (Extended Data Fig. 4b), but display no significant structural similarities to other proteins. The D4 domains play important roles in organizing the intramolecular assembly by interacting with all the other four domains in both the α- and β-subunits (Extended Data Fig. 1i, j). D5_α_ is responsible for interacting with the γ-subunit as described below, whereas D5_β_ mediates homodimerization of the β-subunits.

A β_4_ homotetramer lies at the heart of the α_4_β_4_γ_4_δ_4_ hexadecamer ^7,25^. The β_4_ homotetramer is a dimer of dimers, with each β-subunit interacting with two other molecules (Fig. 2d). The larger dimer interface (β1/β4, and also the β2/β3 counterpart) buries 1,530 Å^2^ surface area in each molecule and involves D5_β_. Both the α- and β-subunits contain a C-terminal CaaX (C, Cys; a, aliphatic residue; X, any residue) prenylation motif, and the corresponding Cys in the α- and β- subunits of rabbit muscle PhK are farnesylated ^26,27^. Mass spectrometry analyses suggest that Cys1220_α_ and Cys1090_β_ in the human muscle PhK are also farnesylated (Extended Data Fig. 4c, d); and interpretable densities are also present for these farnesyl groups (Extended Data Fig. 2g, h). The farnesyl groups on Cys1090_β_ are buried in the larger dimer interface and glue two D5_β_ domains together (Fig. 2d). The smaller β1/β2 (and β3/β4) interface conceals 800 Å^2^ surface area from each subunit and involves regions in D1_β_ and D2_β_. These two β-subunits further interact with two α-subunits, leading to the formation of an α_2_β_2_ heterotetramer.

The α_2_β_2_ heterotetramer also features two different types of α/β interactions (Fig. 2e). The α1/β1 dimer (and also the α2/β2 counterpart) exhibits a pseudo two-fold rotation symmetry, and mainly involves residues in D1_α_ and D1_β_, in particular the Asp68_α_–Arg94_α_ and the Arg110_β_– Arg131_β_ helices, burying 1,630 Å^2^ surface from each protein. The α1/β2 dimer (and α2/β1) buries 1,090 Å^2^ surface from each protein, involves the D1_α_, D2_α_, and D2_β_, and is dominated by polar interactions.

### Autoinhibition of kinase activity

The γ-subunit features a KD that can be further divided into the N- and C-lobes, and a CRD that contains five helices (αJ–αN, Fig. 3a, Extended Data Fig. 5a). The KD structure of human muscle PhK is highly similar to the crystal structure of rPhKγ-KD ^13,14^. Notably, Leu78_γ_, Leu90_γ_, His148_γ_, and Phe169_γ_ line up and form an intact “regulatory spine” (Extended Data Fig. 5b), demonstrating that the KD adopts an active conformation ^28^. The CRD wraps around the kinase C-lobe. The αJ helix interacts with the δ-subunit/calmodulin as described below, whereas the remaining CRD (referred to as <u>a</u>utoinhibitory <u>d</u>omain or AID hereafter) loops back and inserts between the C-lobe and α-subunit.

**Fig. 3.**
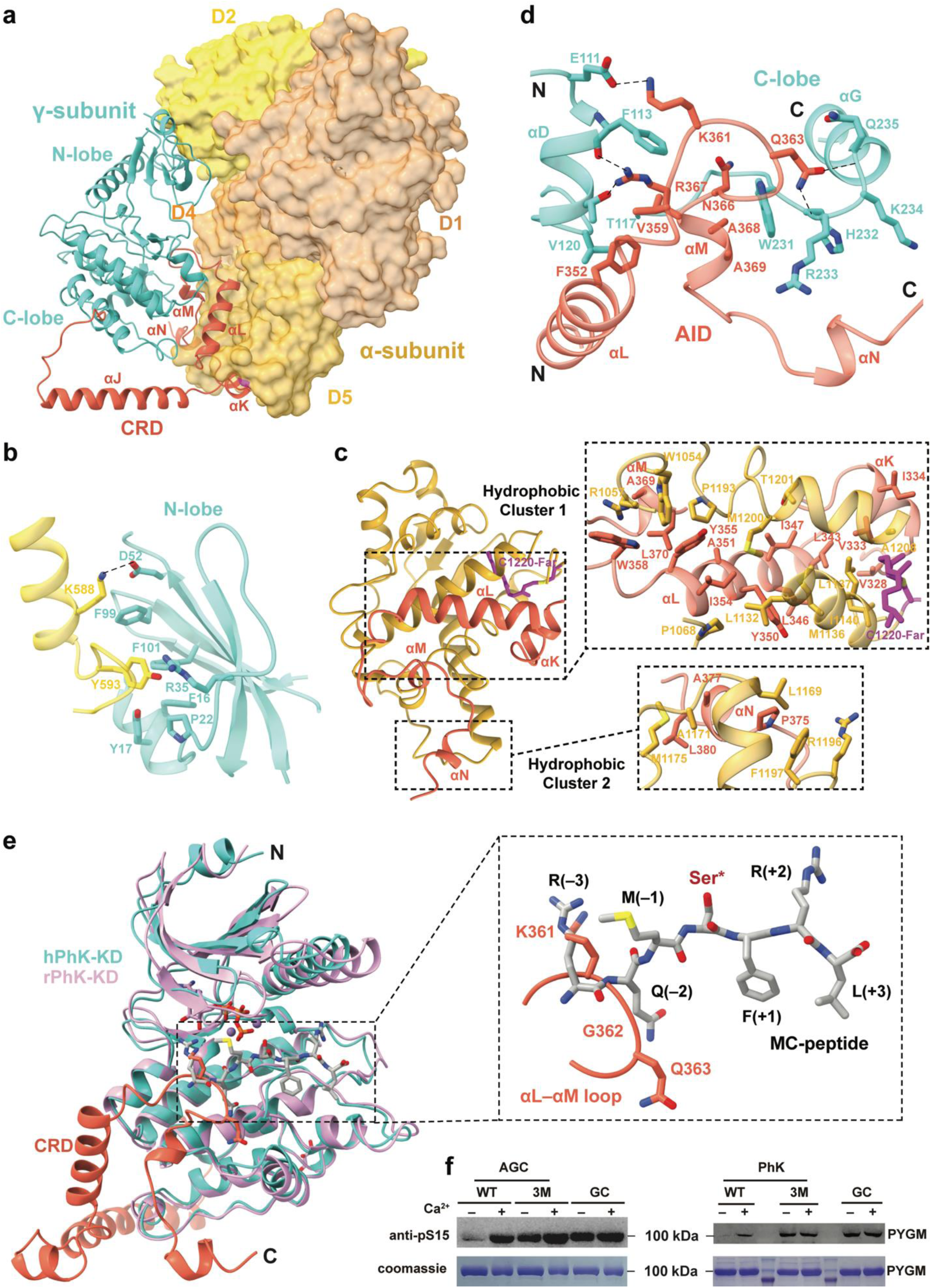
Auto-inhibition of the γ-subunit. **a.** Overall structure of the αγ subcomplex. The α-subunit is shown in a surface representation, whereas the γ-subunit is in ribbons. CRD in the γ-subunit is highlighted in red. **b.** Interactions between the kinase N-lobe of the γ-subunit and the α-subunit. Dash lines indicated polar interactions. **c.** Interactions between the AID of γ-subunit and the α-subunit. **d.** Interactions between AID and the kinase C-lobe. **e.** Structural overlay of hPhKγ and rPhKγ (PDB ID: 1PHK). Lys361_γ_ and Gly362_γ_ occupy the –3 and –2 sites of the substrate peptide in rPhKγ. **f.** The AGC subcomplex and PhK holoenzyme containing the 3M mutant of the γ-subunit, as well as GC, all display Ca^2+^-independent kinase activity.

The γ-subunit is attached to the α-subunit; and contrary to previous models, the β- and γ-subunits do not make contact with each other (Fig. 3a, Fig. 1e). Both KD and AID contribute to interacting with the α-subunit, yielding a large 2,590 Å^2^ binding interface between the two proteins. In the KD, N-lobe residues Phe16_γ_, Tyr17_γ_, Pro22_γ_, Arg35_γ_, Phe99_γ_, and Phe101_γ_ together accommodate Tyr593_α_ in D2_α_; whereas Asp52_γ_ forms an ion pair with Lys588_α_ (Fig. 3b). More extensive interactions are present between AID and the α-subunit (Fig. 3c). Two hydrophobic clusters are present on AID: the larger one involves Val328_γ_, Val333_γ_, Ile334_γ_, Leu343_γ_, Leu346_γ_, Ile347_γ_, Tyr350_γ_, Ala351_γ_, Ile354_γ_, Tyr355_γ_, Trp358_γ_, Ala369_γ_, Leu370_γ_; and the smaller one involves Pro375_γ_, Ala377_γ_, and Leu380_γ_. These hydrophobic residues interact with a range of hydrophobic residues in D5_α_. The farnesyl group on Cys1220_α_ is also involved in this hydrophobic interface. In addition, >20 salt bridge and hydrogen bond interactions are formed between AID and the α-subunit. Together, these extensive interactions firmly anchor the AID onto the α-subunit.

Importantly, the AID also binds to the kinase C-lobe (Fig. 3d). The αL–αM region interacts with the αD helix, as well as the αF–αG loop. This is highly concordant with a previous mass spectrometry study showing that these regions are not surface exposed ^27^. Specifically, Phe352_γ_ and Val359_γ_ cluster with Phe113_γ_, Thr117_γ_, and Val120_γ_. Lys361_γ_ packs against Phe113_γ_, and also forms a salt bridge with Glu111_γ_. Gln363_γ_ contacts His232_γ_–Gln235_γ_. Asn366_γ_, Ala368_γ_, and Ala369_γ_ pack on the Trp231_γ_, whereas Arg367_γ_ contacts Phe113_γ_ and Thr117_γ_. As a result of these interactions, Lys361_γ_–Gly362_γ_ in the αL–αM loop are positioned in such a way that they occupy the –3 and –2 sites of the substrate peptide ^14^ (Fig. 3e). Thus, the AID would inhibit the KD activity by competitively blocking the binding of the substrate. Notably, this pseudo-substrate mechanism highly resembles the autoinhibition mechanism seen in other Ca^2+^/calmodulin-dependent protein kinases (CAMKs) ^29,30^.

The truncated γ-subunit lacking the CRD is constitutively active ^16,17^, demonstrating the autoinhibitory function of this region. To further establish the autoinhibitory role is attributed to the AID, we generated a truncation mutant of the γ-subunit consisting of residues 1–326, and prepared its complex with calmodulin (GC, Extended Data Fig. 3b). Indeed, GC displays Ca^2+^- independent kinase activity towards human muscle GP (PYGM, Fig. 3f). We further designed a triple mutant of the γ-subunit (K361A/Q363A/R367A, 3M) to disrupt the interaction between the αL–αM pseudo-substrate loop and KD (Fig. 3d), and prepared the AGC subcomplex and PhK holoenzyme accordingly. In contrast to wildtype (WT) AGC and PhK, which requires Ca^2+^ for kinase activity, AGC-3M and PhK-3M exhibit constitutive activity towards PYGM regardless of the presence of Ca^2+^ (Fig. 3f). Together, these data validate our structural analyses, and corroborate the critical roles of AID in autoinhibiting the γ-subunit kinase activity.

### A compact conformation of calmodulin

Calmodulin functions in diverse cellular processes and can versatilely bind to hundreds of proteins ^31^. A major difference between PhK and other CAMKs is that calmodulin binds to the γ- subunit tightly even in the absence of Ca^2+^ ion ^11^. In fact, dissociation between the γ-subunit and calmodulin can only be achieved by protein denaturation ^32^. Previous studies suggested two potential calmodulin-binding motifs in the CRD of the rabbit muscle PhK γ-subunit, namely the PhK13 and PhK5 peptides ^15^, which correspond to residues 303–327 and 343–367 in the human γ-subunit, respectively (Extended Data Fig. 5a). Despite the fact that the PhK5 peptide has been involved in binding Ca^2+^/calmodulin ^15,33,34^, our structure unambiguously demonstrates that this region is involved in interacting with the α-subunit and KD (Fig. 3a). Instead, the PhK13 peptide, which constitutes the αJ helix, plays an essential role in interacting with calmodulin (Fig. 4a, Extended Data Fig. 5c). This is consistent with previous crosslinking and hydrogen-deuterium exchange studies ^10,35^.

**Fig. 4.**
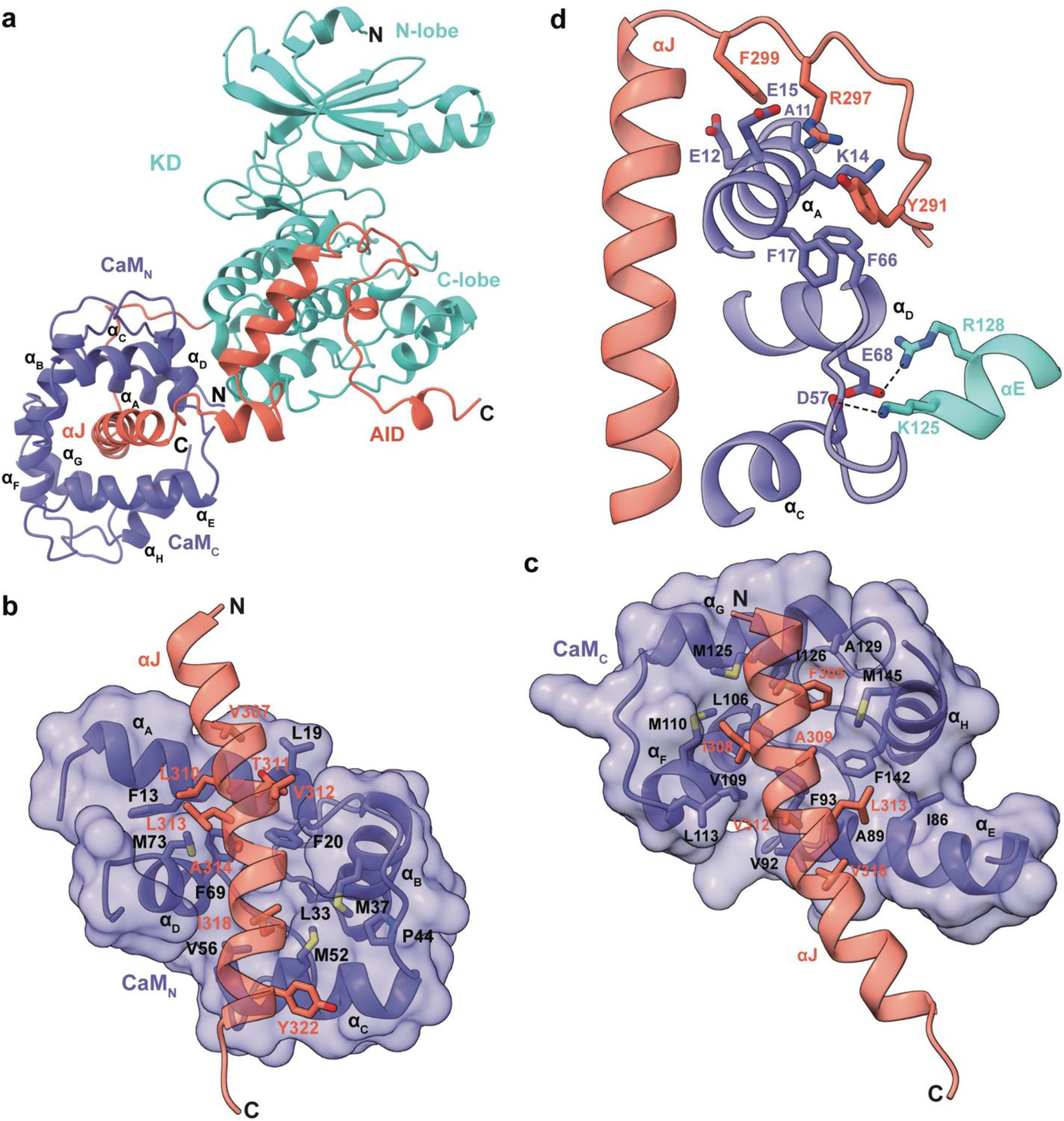
Structure of the γδ subcomplex. **a.** Overall structure of the γδ subcomplex shown in ribbons. The eight helices in calmodulin are labeled α_A_–α_H_. **b.** Interactions between αJ and the N-terminal half of calmodulin. **c.** Interactions between αJ and the C-terminal half of calmodulin. **d.** Interactions between the kinase C-lobe and calmodulin.

Calmodulin adopts a compact conformation, and is firmly held between the αJ helix and the kinase C-lobe. It assumes an ‘antiparallel’ wrapping around the αJ helix, with the C- and N-termini of αJ anchored to the N- and C-terminal halves of calmodulin, respectively (Fig. 4b, c). Calmodulin encompasses the majority of the αJ helix. Over half of the αJ residues are hydrophobic and aromatic, allowing for extensive interactions with the inner hydrophobic surface of calmodulin. Notably, calmodulin also engages with the kinase C-lobe (Fig. 4d). Specifically, Lys125_γ_ and Arg128_γ_ in the αE helix of the kinase C-lobe form bonds with Asp57_δ_ and Glu68_δ_ in the second EF-hand motif of calmodulin. Since these two residues are critical for binding to Ca^2+^, these interactions may offer an explanation for the previous observation that only three Ca^2+^ bind per δ-subunit in PhK ^36^. Moreover, Tyr291_γ_, Arg297_γ_, and Phe299_γ_ in the C-lobe–αJ linker contribute to the interaction with calmodulin by interacting with Ala11_δ_, Glu12_δ_, Lys14_δ_, Glu15_δ_, Phe17_δ_, and Phe66_δ_. Together, these extensive interactions effectively tether calmodulin to the γ-subunit.

### Activation of PhK

To investigate the mechanism of Ca^2+^-induced activation of PhK, we analyzed the cryo-EM structure of PhK in the presence of Ca^2+^ (Extended Data Fig. 2, Table 1). When compared to the inactive state structure, the most notable difference is that the KD–αJ region of the γ-subunit and calmodulin were no longer visible (Fig. 5a). Since the protein sample was intact (Fig. 1b), these regions likely displayed conformational flexibility and were not discernible after particle averaging. The majority of the AID can still be clearly observed, which remain tightly bound to the α-subunit, except for residues 360–365 that harbor the Lys361_γ_–Gly362_γ_ autoinhibitory sites. Ca^2+^-free calmodulin exists in a compact conformation in the inactive PhK structure (Fig. 4a). In contrast, a small-angle scattering study suggested that Ca^2+^/calmodulin adopts an extended conformation when bound to the PhK13 peptide ^33^. It is conceivable that this Ca^2+^-triggered conformational transition of calmodulin would propagate to the γ-subunit via the tight interaction between these two proteins. The KD was likely detached from the α-subunit–AID platform as a result of this conformational change (Fig. 5b), providing an explanation for its absence in the density map. Importantly, the separation of KD from AID would lead to kinase de-inhibition and activation.

**Fig. 5.**
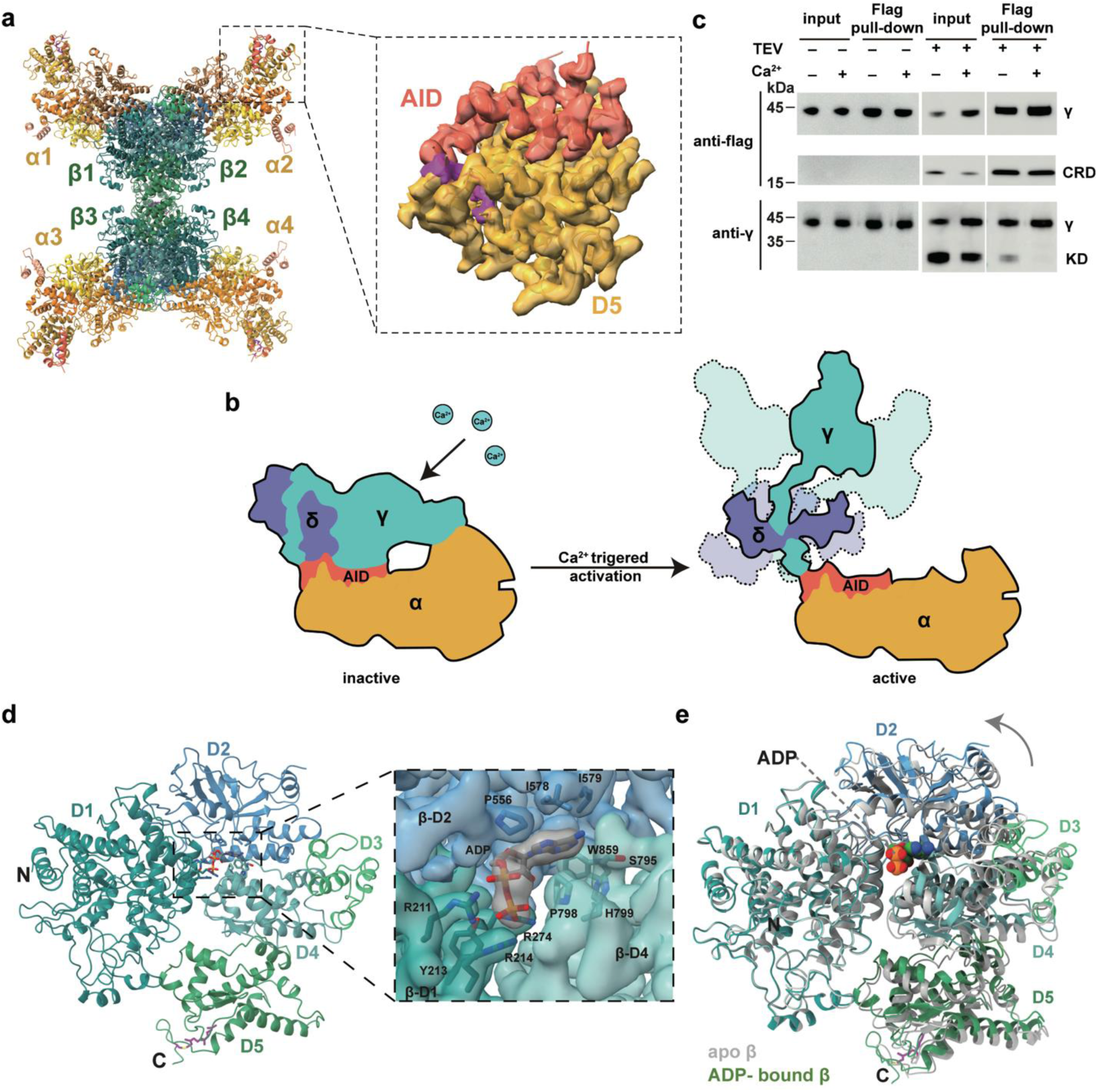
PhK activation. **a.** Cryo-EM structure of PhK in the presence of Ca^2+^. The density map of the D5_α_–AID region is shown on the right. **b.** A model of Ca^2+^-triggered PhK de-inhibition. **c.** After TEV cleavage, significant less amount of KD remains associated with the AG_TEV_C complex upon Ca^2+^ treatment. **d.** Overall structure of the ADP-bound β-subunit. The cryo-EM density map of the ADP-binding pocket is shown on the right. **e.** Structural overlay of the apo and ADP-bound β-subunits.

To biochemically test this scenario, we introduced a tobacco etch virus (TEV) protease cleavage site before CRD in the γ-subunit (Extended Data Fig. 5a), and generated the AG_TEV_C complex. A Flag tag was also engineered onto the C-terminus of the γ-subunit, and was used to perform a pull-down experiment. We reasoned that in the absence of Ca^2+^, the KD would be pulled down by the Flag antibody after it is separated from the CRD by the TEV protease, due to its interactions with AID and the α-subunit (Fig. 3a). In contrast, significant less KD should remain associated upon Ca^2+^ treatment, which leads to calmodulin conformational change and detachment of KD as described above. This hypothesis was borne out by the experiment (Fig. 5c). Together, the structural and biochemical data suggest that calmodulin likely regulates PhK activity using a spring-loaded mechanism: Ca^2+^-free calmodulin binds to the γ-subunit in a compact conformation like a compressed spring; upon sensing Ca^2+^, calmodulin undergoes a conformational change and facilitates detachment of KD and therefore de-inhibition of kinase activity (Fig. 5b).

ADP has also been shown to allosterically enhances PhK activity by binding to the β-subunit ^37^. Indeed, the active PhK structure clearly reveals the presence of ADP bound to the β- subunits, which can be attributed to the ADP present in the sample buffer. Within the structure, ADP is nestled between the D1_β_, D2_β_, and D4_β_ domains (Fig. 5d). The diphosphate moiety is surrounded by Arg211_β_, Tyr213_β_, Arg214_β_, and Arg274_β_, which effectively neutralize the negative charges of the phosphate groups and allow the binding of ADP without divalent metal ions. The presence of ADP leads to a slight conformational change of D2_β_–D5_β_ relative to D1_β_ (Fig. 5e). Notably, D2_β_ interacts with D2_α_ to establish the α1/β2 and α2/β1 dimers (Fig. 2e); whereas D2_α_ interacts with the KD N-lobe and contributes to the maintenance of the inactive state of the γ-subunit (Fig. 3a). The ADP-induced conformational changes in the β-subunits would propagate to the α-subunits through the D2_β_–D2_α_ interaction, facilitating the disruption of the α-subunit–KD interaction and thereby allosterically enhancing PhK activity.

In summary, we have established a structural framework for mechanistic understanding of PhK. We provide the first glance at the structures of the α- and β-subunits, and reveal that the autoinhibition of the γ-subunit kinase activity is achieved through a pseudo-substrate mechanism. Ca^2+^-induced PhK activation is likely achieved by the release of AID-mediated KD autoinhibition as a result of Ca^2+^-dependent conformational change of calmodulin. These findings not only resolve long-standing questions in PhK biochemistry but also lay the foundation for future studies on this important kinase complex.

## Materials and Methods

### Plasmids

The DNA fragments encoding full-length PhK α-, β-, γ-subunits and calmodulin were cloned into pMLink vector (a kind gift from Professor N. Gao, Peking University) for expression in HEK293F cells. A twin-strep tag is engineered to the N-terminal region of the β-subunit. These DNA fragments were then linked into one plasmid following a previously described procedure^38^. Briefly, 1 μg of one plasmid was digested overnight with 10 units PacI (New England Biolabs) in 20 μL at 37 °C, while the other was digested at 25 °C with 10 units SwaI (New England Biolabs). Enzymes were inactivated at 65 °C for 20 min. The PacI digest was treated with 1 units T4 DNA polymerase (New England Biolabs) and 2 mM dCTP, and the SwaI digest was treated with 1 units T4 DNA polymerase and 2 mM dGTP, respectively, both at 12 °C for 15 min. EDTA was added to theses mixtures to a final concentration of 10 mM, before heat inactivation was performed at 75 °C for 20 min. Finally, the two plasmids were mixed and heated to 65 °C and cooled to room temperature for annealing, followed by *E. coli* transformation. Mutations were introduced into the expression plasmids by a PCR-based method. The DNA fragment encoding residues 1–326 of the γ-subunit was cloned into a modified pQLink vector with a N-terminal GST tag and TEV protease cleavage site for expression in *E. coli*. The DNA fragment encoding full-length calmodulin was cloned into a modified pQLink vector with a N-terminal 8 ×His tag and TEV cleavage site. These two genes were linked into one plasmid following the procedure described above. The DNA fragment encoding full-length human muscle phosphorylase (PYGM) were cloned into a modified pCS2 vector with an N-terminal twin-strep tag and TEV cleavage site. The DNA fragment encoding human lysosomal α-glucosidase (GAA) residue 70-952 was cloned into the pMLink vector, and engineered with an N-terminal IL-2 signal peptide and 8 ×His tag.

### Protein expression and purification in HEK293F cells

HEK293F cells were cultured in SMM 293T-I medium (Sino Biological Inc.) at 37 °C, with 5% CO_2_ and 55% humidity. The plasmid expressing PhK complex was transfected into the cells using polyethylenimine (PEI, Polysciences). After 36 hours of transfection, the cells were collected by centrifugation at 500 ×g and resuspended in buffer A [50 mM HEPES, pH 6.8, 100 mM NaCl, 30 mM β-glycerolphosphate, 10% w/v sucrose and 2 mM dithiothreitol (DTT)], supplemented with Protease Inhibitor Cocktail (Bimake). The cells were then disrupted by sonication. The insoluble debris was removed by centrifugation, and the resulting supernatant was incubated with the StrepTactin beads (Smart Lifesciences) at 4 °C for 1 h with gentle shaking. The beads were washed with buffer A supplemented with 10 mM EDTA and eluted using buffer A supplemented with 5 mM desthiobiotin (IBA). Next, the purified protein was treated with the λ-phosphatase in the presence of 2 mM MnCl_2_ at room temperature for 1 hour. The PhK mutant and the αγδ subcomplexes were purified similarly.

For the purification of PYGM, the cell lysate was incubated with the StrepTactin beads in buffer B (50 mM Tris-HCl, pH 8.0, 500 mM NaCl, 10 % v/v glycerol, and 2 mM DTT), supplemented with 1 mM PMSF (phenylmethylsulfonyl fluoride). The beads were washed with buffer B and the elution was performed using buffer B supplemented with 5 mM desthiobiotin. The eluted protein was digested with the TEV protease overnight and further purified by gel filtration chromatography using a Superose 6 column (GE Healthcare), eluted in buffer C (25 mM HEPES, pH 7.5, 150 mM NaCl and 2 mM DTT).

For the purification of GAA, the plasmid was transfected into the HEK293F cells using PEI. The conditioned medium was collected 4 days after transfection, buffer-exchanged into the binding buffer (25 mM Tris-HCl, pH 7.4, 150 mM NaCl), and then incubated with Ni-NTA resin at 4 °C for 1 h. The beads were subsequently washed with the binding supplemented with 20 mM imidazole, and eluted by the binding buffer supplemented with 300 mM imidazole.

### Protein expression and purification in *E. coli*

The *E. coli* BL21(DE3) bacterial cultures were grown at 37 °C in the LB (Luria-Bertani) medium until reaching an OD_600_ of 0.6–0.8 before induced with 0.5 mM isopropyl β-D-1- thiogalactopyranoside (IPTG) at 18 °C for 20 h. The cells were collected by centrifugation at 6,000 rpm and were resuspended in buffer D (50 mM Tris-HCl, pH 8.0, 500 mM NaCl, and 5 mM β-mercaptoethanol). The cells were then disrupted by sonication in the presence of 1 mM PMSF. Following sonication, the insoluble debris was removed by centrifugation. For the purification of G_326_C complex, the supernatant was incubated with the glutathione beads at 4 °C for 1 hour with gentle shaking, and the recombinant protein was eluted using buffer D supplemented with 10 mM reduced L-glutathione (Sigma–Aldrich). The purified protein was digested with TEV protease overnight. The untagged protein was further purified by gel filtration chromatography using a Superdex 200 10/300 GL column (GE Healthcare), eluted in buffer E (25 mM HEPES, pH 6.8, 150 mM NaCl, and 2 mM DTT). The purified protein was then incubated with glutathione beads to remove the GST fusion tag, followed by another gel filtration chromatography step and eluted using buffer E. For the purification of calmodulin, the cell lysates in buffer D was incubated with Ni-NTA resin (GE Healthcare), washed using buffer D supplemented with 20 mM imidazole, and eluted using buffer D supplemented with 300 mM imidazole. The purified calmodulin was digested with TEV protease, and the untagged calmodulin was further purified by gel filtration chromatography using a Superdex 200 10/300 GL column, eluted in buffer C.

### Cryo-electron microscopy sample preparation

For preparing the cryo-EM sample of inactive PhK, dephosphorylated PhK was incubated with purified calmodulin using 1:20 molar ratio in the presence of 4 mM ethylene glycol-bis (β-aminoethyl ether)-tetraacetic acid (EGTA), and further purified using a Superose 6 column, eluted in buffer F (25 mM HEPES, pH 6.8, 150 mM NaCl, and 2 mM DTT). Prior to cryo-EM sample preparation, 4 mM EGTA and additional calmodulin (4-fold excess) were added to the protein sample, to ensure that calmodulin does not fall off. For the sample of Ca^2+^-activated PhK, the dephosphorylated protein was first incubated with excessive calmodulin as described above, in the presence of 1 mM CaCl_2_ on ice for 1 hour, and then purified by Superose 6 column, eluted in buffer G (25 mM HEPES, pH 8.2, 150 mM NaCl, 1 mM CaCl_2_, and 2 mM DTT). Prior to cryo-EM sample preparation, additional calmodulin (4-fold excess) was added, and the mixture was incubated on ice for 1 hour, in the presence of 1 mM CaCl_2_, 1 mM ADP, 1 mM AlCl_3_, 4.5 mM NaF, 2 mM MgCl_2_.

Holey carbon grids (Quantifoil R1.2/1.3) were coated by continues carbon film using mica plate and glow-discharged for 30 seconds using a plasma cleaner (Harrick PDC-32G-2). The sample preparation was carried out using a Vitrobot Mark IV (FEI). Specifically, 4 μL aliquots of the PhK holoenzyme (0.45 mg/ml) were applied onto the grids, allowing for a 5-second incubation at 4 °C and 100% humidity. Subsequently, the grids were blotted with filter paper (Tedpella) for 0.5 seconds using a blotting force of –1, and immediately plunged into liquid ethane. Grids screening was performed using a 200 kV Talos Arctica microscope equipped with Ceta camera (FEI). Good grids were transferred to a 300 kV Titan Krios electron microscope (FEI) for data collection.

Data were acquired using the EPU software (E Pluribus Unum, Thermo Fisher) on a K3 Summit direct electron detector (Gatan) operating in a super-resolution mode. The defocus range was set from –1 to –1.5 μm. Micrographs were recorded at a nominal magnification of 81,000 (pixel size of 1.07 Å at the object scale). The micrographs were dose-fractioned into 40 frames with dose rate of around 21.47 electrons per pixel per second for a total exposure time of 3.2 s.

### Imaging processing

The data were processed using cryoSPARC (version 3)^39^. Movie stacks were collected and motion-corrected using the patch motion correction, and the CTF parameters were determined using the patch CTF estimation. Summed images were then manually screened to remove low-quality images through exposure curation. Initially, particle picking was performed using a blob picker, and templates were generated through subsequent 2D classification. The particles from template picking were subjected to multiple rounds of 2D classification to exclude inaccurate particles. The selected particles were then subjected to ab-initio reconstruction and heterogeneous refinement to select the appropriate particles. The particles resulting from the heterogeneous refinement were then utilized in homogeneous refinement, ultimately leading to the generation of the final 3D reconstruction. To obtain higher-quality local density map, mask-based local refinement was further performed using cryoSPARC. All maps were then sharpened using DeepEMhancer^40^. The local resolution map was produced with the local resolution estimation program in cryoSPARC and displayed using UCSF Chimera^41^.

### Model building and structure refinement

Initial models of PhK α, β subunit and the KD of γ subunits were generated using AlphaFold2^42,43^, and calmodulin were obtained from a previous crystal structure (PDB ID: 1CDL)^44^. These models were docked into the cryo-EM density map using UCSF Chimera. The CRD of the γ-subunit was built *de novo* using Coot^45^. Further structure model building was performed using Coot and refined using the real-space refinement in Phenix^46^. Figures were prepared with UCSF ChimeraX ^47^.

### Glucoamylase activity assay

To determine glucoamylase activity, 100 nM PhK holoenzyme or 400 nM αγδ subcomplex was incubated with 15 mg/mL maltose (Macklin), isomaltose (Macklin) or glycogen (from bovine liver, Sigma–Aldrich) at 37 °C for 1 h in the reaction buffer containing 50 mM HEPES, pH 6.8, 100 mM NaCl. The activity was measured by quantifying the amount of released glucose using Glucose Assay Kit (Sigma–Aldrich) in 96-well plate. GAA was used as a positive control, and the reaction buffer for GAA is containing 50 mM sodium acetate, pH 4.5, 100 mM NaCl.

### Kinase assay

A concentration of 100 nM of PhK, PhK_3M_ and G_326_C, or 40 nM of AGC subcomplex, AG_3M_C and G_326_C were incubated with 2 µM PYGM in a reaction buffer containing 25 mM HEPES, 100 mM NaCl, 1 mM ATP, 5 mM MgCl_2_, and either 0.5 mM EGTA at pH 6.8 or 1 mM CaCl_2_ at pH 8.2. The assays were carried out at 30 °C for 10 min and terminated by the addition of SDS-PAGE sample buffer (TransGen Biotech) supplemented with 20 mM EDTA, and then boiled. The samples were then resolved by SDS-PAGE and analyzed by immunoblotting using the anti-PYGL (phospho S15) antibody (Abcam, ab227043).

### TEV cleavage and FLAG pull-down assay

The PhK γ-subunit was modified by introducing a N-terminal 6 ×his tag and a C-terminal FLAG tag. For cleavage, the sequence of TEV protease was introduced into the γ-subunit replacing residue 293–299. The AG_TEV_C complex was purified following the procedure described above. For the pull-down experiments, the AG_TEV_C complex (0.05 mg/mL) was first incubated with either 0.5 mM EGTA or 0.5 mM CaCl2 at 4 °C for 15 min. Subsequently, the AG_TEV_C complex, excess TEV protease and the FLAG beads (Smart Lifesciences) were incubated at 4 °C for 1 h with gentle shaking in the binding buffers (25 mM HEPES, pH 6.8, 150 mM NaCl, 2 mM DTT, supplemented with either 0.5 mM EGTA or 0.5 mM CaCl_2_). The beads were centrifuged at 500 × g and then washed three times with the binding buffers. The proteins bound to the beads were eluted using the binding buffers supplemented with 0.2 mg/mL 3 × FLAG peptide (NJPeptide). The eluted proteins were then analyzed using SDS-PAGE and visualized through Coomassie blue staining or immunoblotting using antibodies for the *PhKG1* (Abcam, ab194112) and FLAG tag (Sigma, F3165).

## Acknowledgments

We thank the Core Facilities at the School of Life Sciences, Peking University for help with negative-staining EM; the Cryo-EM Platform of Peking University for help with data collection; and the High-performance Computing Platform of Peking University for help with computation. We also thank the National Center for Protein Sciences at Peking University for assistance with the BioTek Cytation Reader and mass spectrometry facilities. The work was partly supported by the Qidong-SLS Innovation Fund to J.X. and by Changping Laboratory.

## Author Contributions

XY and MZ contributed equally to this work. XY and MZ carried out the structural and biochemical studies, with YW providing critical assistance. XL performed mass spectrometry analyses. JX conceived and supervised the project. JX and XY wrote the manuscript, with inputs from all authors.

## Data and materials availability

Cryo-EM density maps of PhK have been deposited in the Electron Microscopy Data Bank with accession codes EMD-36212 (inactive, overall), EMD-36214 (inactive, αβγδ subcomplex), EMD-36215 (inactive, γδ subcomplex), EMD-36213 (Ca^2+^, overall), and EMD-36216 (Ca^2+^, αγ subcomplex). Structural coordinates have been deposited in the Protein Data Bank with the accession codes 8JFK and 8JFL.

---

**Extended Data Fig. 1.**
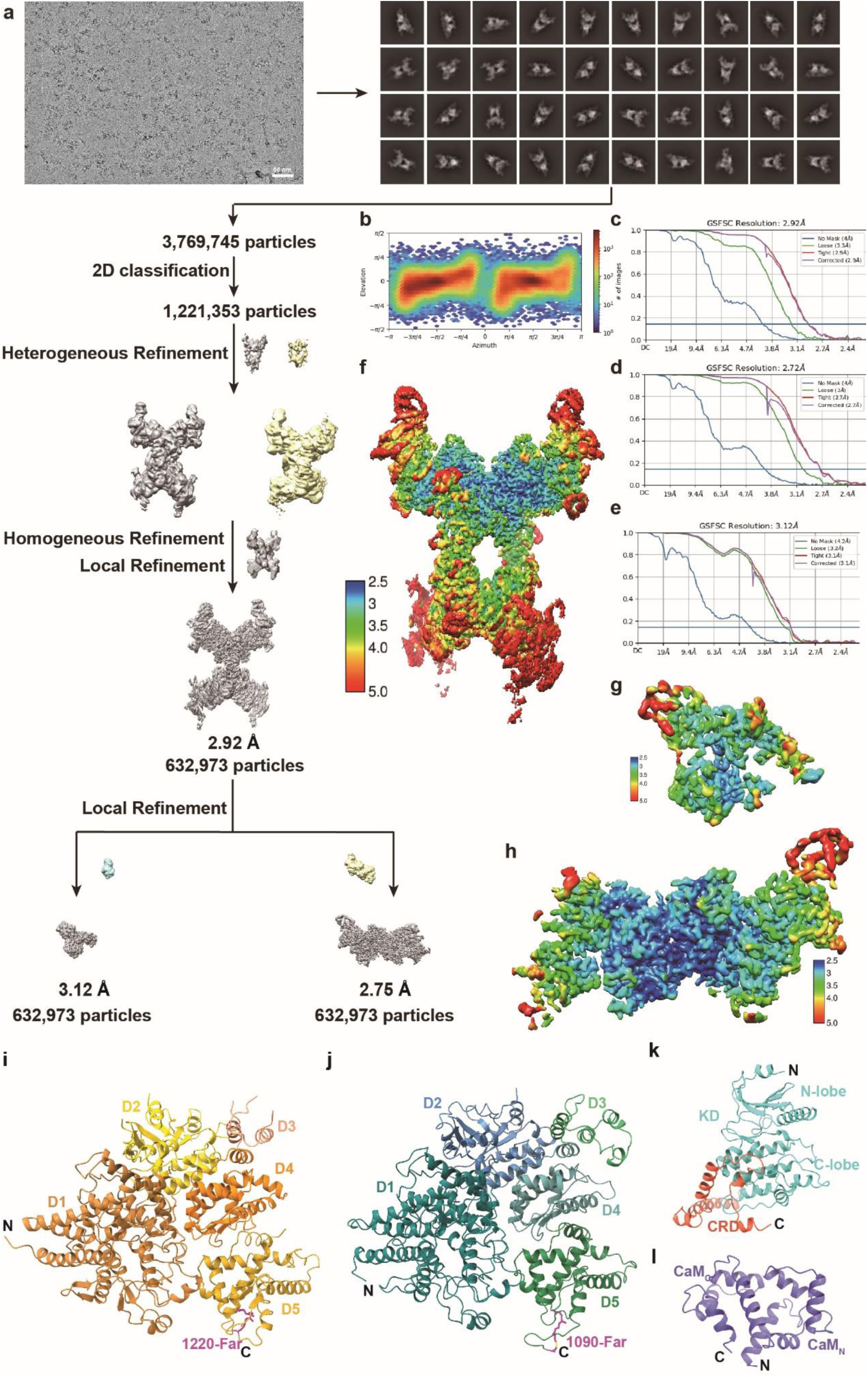
Cryo-EM 3D reconstruction of the inactive PhK complex. **a.** Flowchart of cryo-EM data processing. **b.** Angular particle distribution heat map. **c–e.** Gold-standard Fourier shell correlation (GSFSC) curves of the inactive PhK holoenzyme, αβγδ subcomplex, and γδ subcomplex, respectively. **f–h.** Resolution estimations for the final maps of the inactive hPhK holoenzyme, αβγδ subcomplex, and γδ subcomplex. **i–l.** Individual structures of the α-, β-, γ-, and δ-subunits are shown in ribbons.

**Extended Data Fig. 2.**
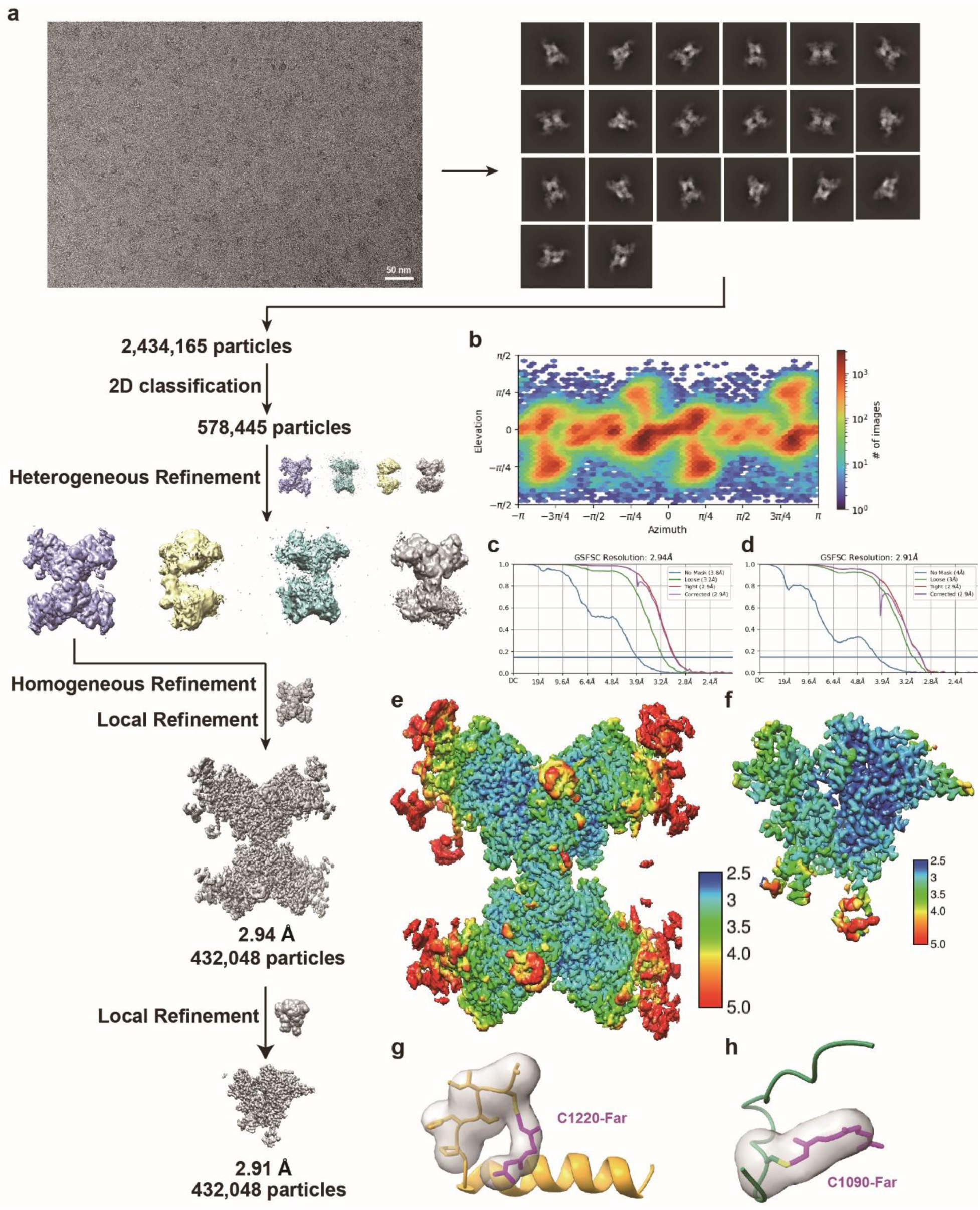
Cryo-EM 3D reconstruction of the active PhK complex. **a.** Flowchart of cryo-EM data processing. **b.** Angular particle distribution heat map. **c, d.** Gold-standard Fourier shell correlation (GSFSC) curves of the active PhK and the αγ subcomplex **e, f.** Resolution estimations for the final maps of the active PhK and the αγ subcomplex. **g.** Density map for the farnesyl group on the α-subunit. **h.** Density map for the farnesyl group on the β-subunit.

**Extended Data Fig. 3.**
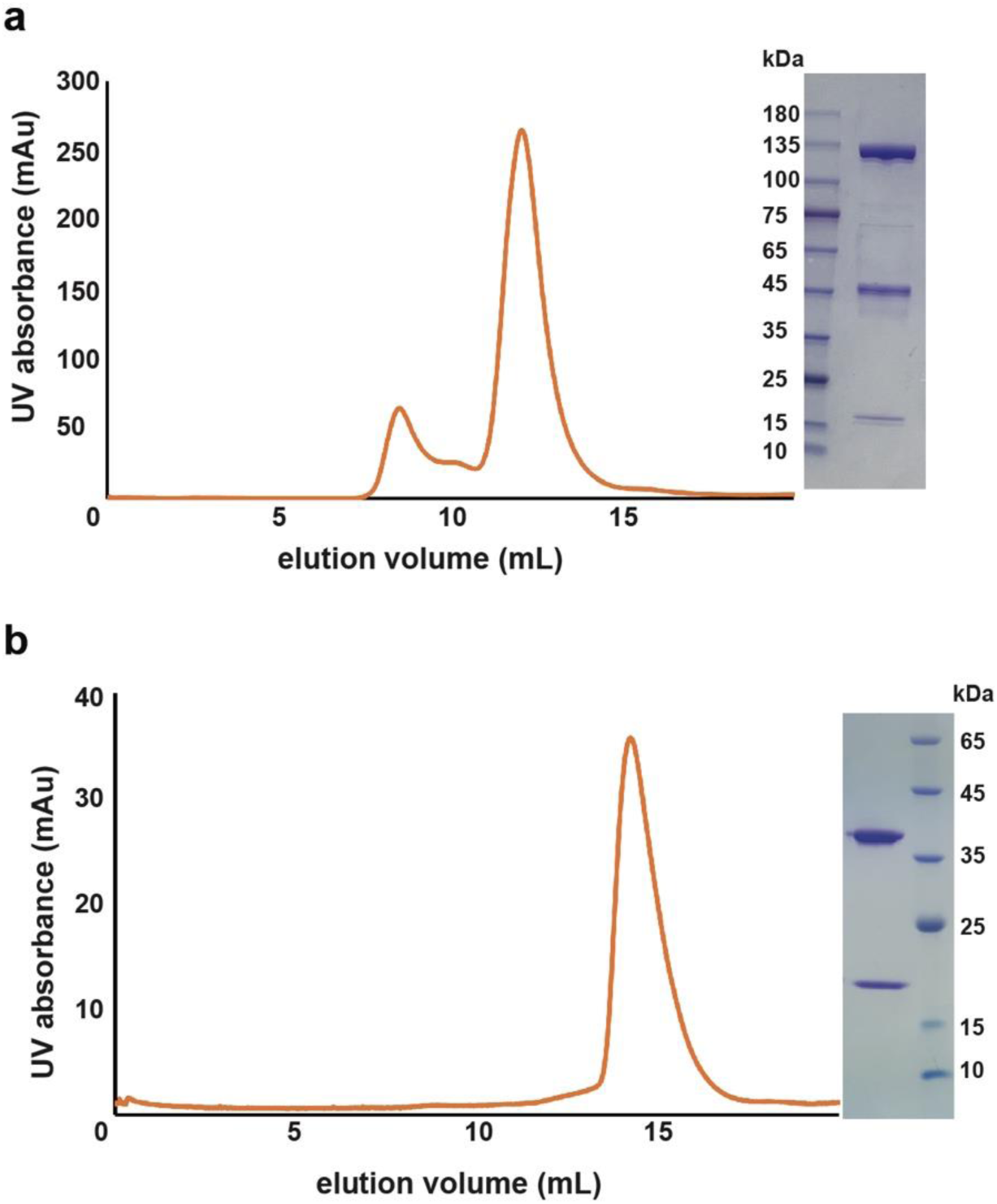
Purification of the AGC and GC subcomplexes. **a.** Size-exclusion chromatography and SDS-PAGE analysis of the AGC complex. **b.** Size-exclusion chromatography and SDS-PAGE analysis of the GC complex.

**Extended Data Fig. 4.**
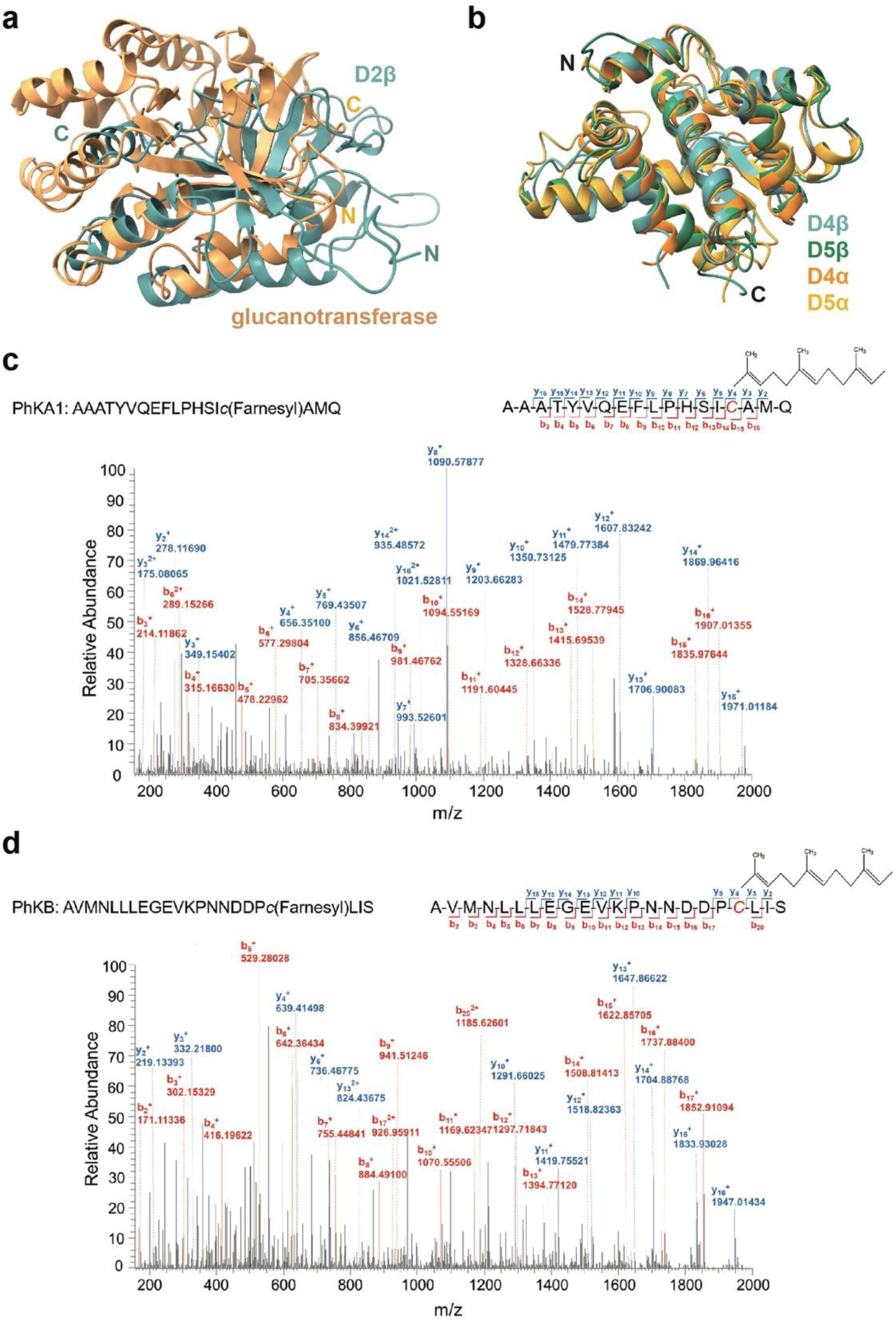
Structures of the α- and β-subunits. **a.** Structural overlay of the D2 domain of the β-subunit and a glucanotransferase (PDB: 1K1Y), shown in cyan and yellow, respectively. **b.** Structural overlay of the D4 and D5 domains of the α- and β-subunits. **c.** Mass spectrometry analysis suggests that Cys1220_α_ in human PhK is farnesylated. **d.** Cys1090_β_ is also farnesylated.

**Extended Data Fig. 5.**
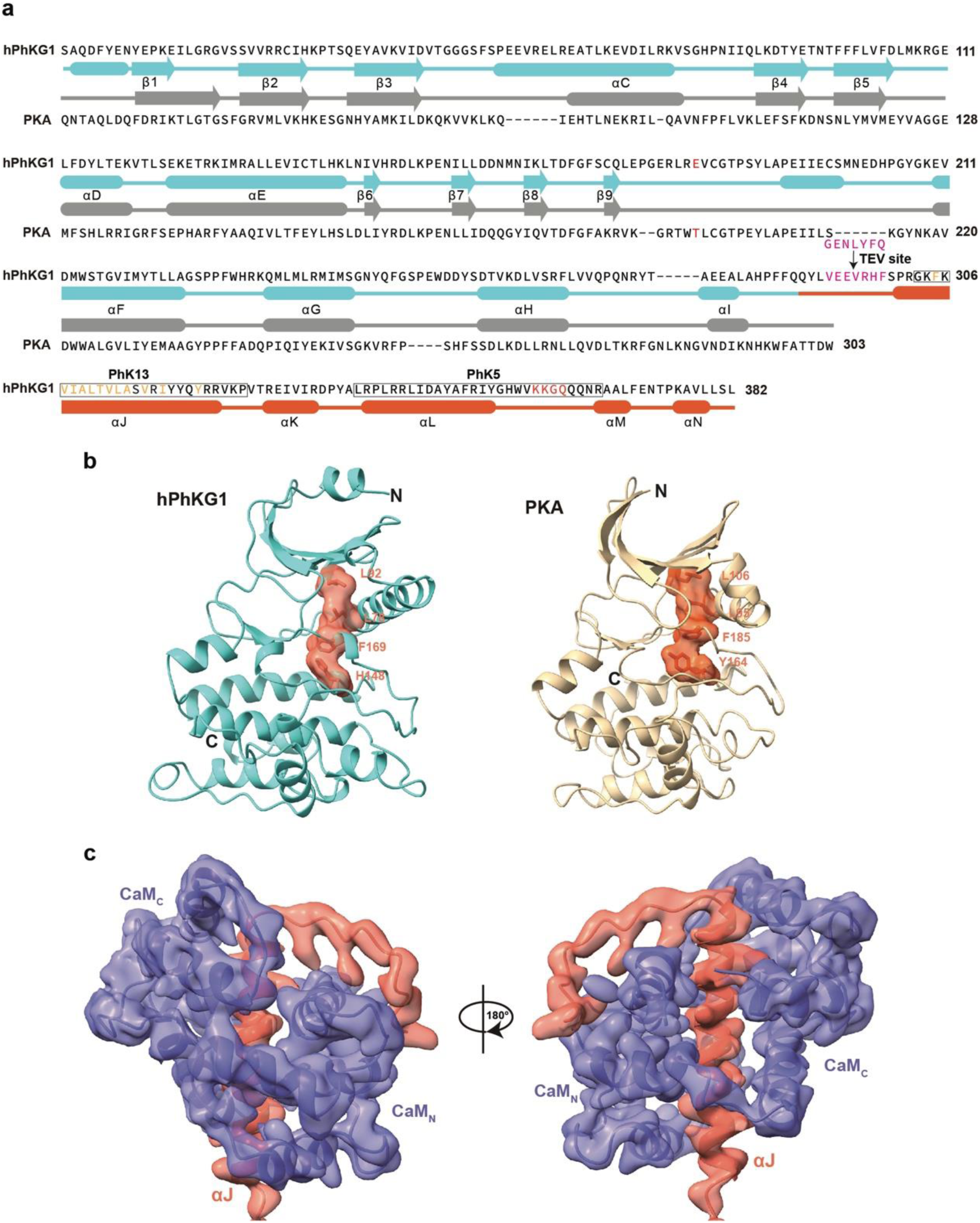
Structure of the γ-subunit and its interaction with calmodulin. **a.** Sequence alignment between the γ-subunit and PKA. The secondary structures of the two proteins are indicated. Glu183_γ_ and Thr197_PKA_ are highlighted in red. Residues 293–299 of PhKγ are replaced by the TEV cleavage site (ENLYFQG) in the AG_TEV_C mutant. The PhK13 and PhK5 peptide regions are highlighted using black boxes. **b.** Leu78_γ_, Leu90_γ_, His148_γ_, and Phe169_γ_form an intact “regulatory spine” in the γ-subunit. **c.** The cryo-EM density map of αJ–calmodulin region shown in two orientations.

